# Hepatic Cholesteryl Ester Transfer Protein Regulates Sex-specific Liver Metabolic Adaptation and Metabolic-Associated Steatotic Liver Disease Risk in Diet-induced Obesity

**DOI:** 10.64898/2026.06.28.735072

**Authors:** Sivaprakasam Chinnarasu, Uche Anozie, Lin Zhu, John M. Stafford

## Abstract

Metabolic dysfunction-Associated Steatotic Liver Disease (MASLD) and associated dyslipidemia is a growing health issue that gives rise to cardiovascular risk. Men are more prone to development of MASLD than women. Understanding mechanisms underlying sex differences in MASLD may lead to improved prevention and treatment approaches. Cholesteryl ester transfer protein (CETP) is a lipid transfer protein that shuttles triglycerides and cholesteryl esters between blood lipoproteins and tissues. In this study investigate the impact of hepatic CETP expression on MASLD. Hepatic CETP expression (L-HuCETP) was achieved by injecting liver-targeted CETP-expressing adeno-associated virus into C57BL/6J mice. In females, L-HuCETP improved glucose tolerance, consistent with our prior clamp results in global human CETP transgenic mice. Whereas in males, L-HuCETP worsened glucose metabolism and impaired insulin signaling. Correspondingly, L-HuCETP expression reduced the expression of gluconeogenic pathway genes in females but upregulated these genes in males. In males, L-HuCETP mice exhibited increased hepatic lipid droplet accumulation, lipogenesis proteins and these changes were not observed in females. L-HuCETP expression resulted in sex-specific hepatic responses, with increased expression of inflammation and fibrosis related genes in male, but decreased expression of these genes in females. Mechanistic studies indicate that L-HuCETP had sex specific effects on transcription factors ChREBP and HNF4α, which are important for glucose and lipid metabolism. Our studies suggest that sex-specific roles of L-HuCETP with regard to liver metabolic adaptation and MASLD risk in obesity, highlighting CETP-mediated pathways as potential targets for sex-specific precision medicine approaches to improve MASLD.

## 1. Introduction

The liver serves as the central hub of lipid metabolism, regulating both circulating and hepatic lipid homeostasis. Hepatic lipid metabolic homeostasis is maintained through several coordinated processes: 1) uptake of dietary lipids *via* lipoprotein receptors; 2) *de novo* lipogenesis; 3) secretion of triglycerides (TG) in Very Low-Density Lipoprotein (VLDL-TG); and 4) fatty acid oxidation and bile acid synthesis and excretion. Any impairment of these coordinated steps may contribute lipid accumulation in hepatocytes and promote metabolic dysfunction-associated steatotic liver disease (MASLD), formerly referred to as nonalcoholic fatty liver disease (NAFLD). MASLD spans a spectrum of liver pathology, starting with simple steatosis and potentially progressing to inflammatory injury, termed Metabolic Dysfunction-Associated Steatohepatitis (MASH), and occasionally, advancing to cirrhosis and hepatocellular carcinoma. MASLD is currently the leading cause of chronic liver disease in both the United States and worldwide [1–3].

It is estimated more than 20-30% of general populations is impacted by MASLD driven by the high prevalence of obesity and insulin resistance. MASLD is more prevalent in men, who are also more likely to develop steatohepatitis, fibrosis and liver related complications compared with women. Several factors such as genetic influences, hormones, and lifestyles contribute to sex differences in MASLD development and progression. Many studies have demonstrated protective effects of estrogenic pathways in MASLD in both humans and animals [4–7]. By contrast, the role of androgens in MASLD risk and progression are not clear, with many studies suggesting protective effects of androgens reviewed in [8]. In men, testosterone deficiency leads to hepatic steatosis and insulin resistance and can be reversed by testosterone replacement therapy [9]. Men with low testosterone levels have exhibited higher visceral adiposity, greater insulin resistance, and an elevated hepatic steatosis index compared with men with higher total testosterone levels [10]. Interestingly, no significant difference in hepatic fibrosis was observed between men with low and high testosterone levels [11]. Testosterone therapy does not increase cardiovascular complications in hypogonadal men compared with placebo [12] although may improve glycemic control or, in some cases, may reverse type 2 diabetes–related complications in high-risk individuals [13]. Notably, Mendelian randomization analyses suggest that genetically predicted higher circulating testosterone levels are associated with increased risk of coronary artery disease and elevated blood pressure [14], highlighting the complex and context-dependent role of androgens in cardiovascular disease for men.

Cholesteryl ester transfer protein (CETP) is a circulating lipid transfer protein which is highly expressed in the liver and adipose tissue. CETP mediates the shuttling of triglycerides (TG) and cholesteryl esters between high-density lipoprotein (HDL) to apolipoprotein B (ApoB)-containing lipoproteins [15]. CETP activity is known to influence HDL-cholesterol (HDL-C) levels, as shown in both preclinical and clinical studies. Early findings have shown that rare genetic CETP deficiency significantly increases the HDL size and number [16]. Mice naturally lack CETP and exhibit high HDL-C levels, which contributes to their lipoprotein profile differing substantially from that of humans and may explain their relative resistance to atherosclerosis. In contrast, humans have high levels of low-density lipoprotein (LDL) cholesterol, an atherogenic lipoprotein fraction. Transgenic expression of CETP in mice produces a more human-like lipoprotein profile, often with CETP activity exceeding that observed in humans. Our previous studies shown that CETP plays a critical role in hepatic lipid metabolism and circulating TG levels [17–19]. In female mice, overexpression of CETP driven by a simian promoter protects against diet induced insulin resistance, enhances liver lipid oxidation and reduces VLDL-TG production [6, 18, 20]. Similarly, transgenic expression of the human CETP (HuCETP) gene driven by its natural flanking sequences improves diet induced fatty liver and insulin resistance in female mice [21]. By contrast to the protective metabolic role of CETP in female mice, expression of CETP in male mice raises plasma VLDL-TG levels by impairing TG clearance through liver androgen receptor-dependent mechanism [22]. Since CETP is largely expressed in the liver in humans, we postulated that hepatic-directed expression of CETP might influence sex specific risk of MASLD.

In the present study, we characterized the contribution of hepatic CETP to diet-induced metabolic phenotypes in male and female mice using adeno associated virus (AAV) mediated, liver specific human CETP expression model (L-HuCETP) compared to AVV-GFP control. We demonstrate that female mice rely on hepatic HuCETP to confer protection against high-fat-diet induced insulin resistance. These protective effects include improved glucose intolerance, decreased expression of gluconeogenic genes, and increased circulating estrogen levels. In contrast, hepatic huCETP expression in obese male mice worsens insulin resistance, accompanied by exacerbated upregulation of gluconeogenic genes, increased hepatic TG accumulation and inflammation, and no detectable changes in circulating hormone levels. These results highlight a sex-specific role for hepatic CETP in regulating metabolic adaptation and susceptibility to MASLD in the context of obesity.

## 2. Material and Methods

### 2.1 Animals and diets

Male and female (C57BL/6J) mice aged 6 weeks were purchased from Jackson laboratory. All mouse experiments were approved by the Vanderbilt University Institutional Animal Care and Use Committee. Mice were housed under controlled temperature and humidity conditions on a 12-hour light/dark cycle with ad-libitum access to a standard chow diet and water. To induce obesity, mice were fed a high fat diet (HFD, 60%kcal fat) (cat # D12492) for 15 weeks.

### 2.2 AAV8-TBG-HuCETP injection

AAV8-TBG-HuCETP and AAV8-TBG-GFP vectors were purchased from Penn Vector at the University of Pennsylvania. 1×10^11^ GC/Kg of vector were injected via retro-orbital route when mice were 8 weeks old. After two weeks of injection, plasma samples were collected and CETP protein expression in mice were confirmed.

### 2.3 Blood lipids, enzyme activities and hormones

Blood samples collected after 5 hours fasting and plasma was isolated for determination of lipids, enzyme activities, and hormone levels. Plasma samples were used to measure cholesterol and triglyceride levels followed by manufacturer’s protocol. Ultra-sensitive Mouse Insulin Elisa kit from crystal chem (cat # 90080) was used to determined insulin concentration following manufacturer’s instructions. AST and ALT activities were assessed using the Cayman chemical activity assay kit (Cat#701640; cat# 700260). CETP activity was measured using a cell-free assay according to the manufacturer’s protocol (Roar Biomedical) as we reported previously[19]. Free fatty acid ((cat# 700310), Estradiol (cat# 501890) and testosterone (cat# 80552) levels were measured according to manufacturer’s (Cayman chemical) instructions.

### 2.4 Plasma lipoprotein separation

To separate plasma lipoproteins, we used size exclusion chromatography (SEC) with a high-performance liquid chromatography (HPLC) system. The SEC separation was performed as described with a superose™ 6 10/300 GL from GE Healthcare (Cat# 17-5172-01). Fractional cholesterol and TG content was assayed using Infinity™ Cholesterol (Cat# TR13421) and TG (Cat# TR22421) reagents.

### 2.5 Oral glucose tolerance test (OGTT)

An oral glucose tolerance test was performed with AAV8-TBG-GFP/HuCETP mice 8 weeks after HFD. Mice were fasted at 7 AM and an OGTT was performed after 5 hours fasting. Mice were administered 20% (w/v) dextrose by oral gavage at a dose of 2 g/kg body weight, and tail-vein blood glucose levels were measured using a glucometer (Accu-Chek, Accu-Chek Aviva Plus Meter) at 10 min before gavage (−10 min) and 5, 10, 15, 30, 45, 60, 90, and 120 min after gavage. We used trapezoidal method to calculate the area under the curve of glucose excursion.

### 2.6 Subcellular isolation

Subcellular isolation was performed according to Strzelecki et al. [23]. Briefly, frozen liver tissue (∼100 mg) was thawed on ice, finely minced, and suspended in 3 mL of homogenization buffer containing 0.33 M sucrose, 10 mM MgCl, and 50 mM Tris–HCl (pH 7.8) supplemented with protease and phosphatase inhibitors. Tissues were homogenized at 0 °C using a Teflon–glass Potter–Elvehjem homogenizer. The homogenate was centrifuged at 100 × g for 10 min at 4 °C to pellet nuclei and unbroken cells. The resulting supernatant was collected as the cytosolic fraction. The nuclear pellet was washed once with PBS and resuspended in nuclear extraction buffer for downstream analyses. Protein concentrations were determined prior to immunoblotting.

### 2.7 Immunoblotting

For the analysis of circulating apolipoproteins, 2 µL of plasma was used for each immunoblotting. For hepatic protein analysis, 30–40 µg of total liver proteins was loaded per lane to determine fatty acid synthase (FASN), acetyl-CoA carboxylase 1 (ACC1), stearoyl-CoA desaturase 1 (SCD1), low-density lipoprotein receptor–related protein 1 (LRP1), and β-actin. Protein expression of Carbohydrate response element binding protein (ChREBP), Hepatocyte nuclear factor 4-alpha (HNF4α), and Forkhead box protein O1 (FOXO1), along with Histone H3 and alpha-tubulin as nuclear and cytosolic markers, respectively, was analyzed using liver cytosolic and nuclear fractions. Detailed antibody information is provided in Supplementary Table1.

### 2.8 RNA isolation and gene expression

Tissue samples were flash-frozen in liquid nitrogen immediately when mice were sacrificed and stored at −80 °C until later use. Liver tissue (25 mg) was bead homogenized, and total RNA was isolated according to manufacturer instructions (RNA Miniprep Kit, Zymo Research, Irvine, CA, USA). For qPCR: complementary DNA was synthesized from 1 μg of RNA (iScript, Bio-Rad, Hercules, CA, USA). Quantitative PCR was performed in duplicates using primers (supplemental Table 1) and reagents.

### 2.9 Histology and Imaging

Liver tissues from the left lobe of each mouse were fixed in 10% neutral-buffered formalin for section preparation. Samples were processed by the Vanderbilt University Translational Pathology Shared Resource (TPSR) Core, where tissues were embedded, sectioned at 5-µm thickness, and stained with picrosirius red and Masson’s trichrome. Stained sections were mounted on glass slides and imaged using a light microscope (AmScope).

### 2.10 Insulin injection via portal vein

Mice maintained on a high-fat diet (HFD) were fasted for 5 hours prior to the experiment. Following the fasting period, mice were anesthetized, and the abdominal cavity was surgically exposed through a U-shaped skin incision. The intestines were gently displaced to provide clear access to the portal vein. A baseline liver tissue sample was collected from the left hepatic lobe before insulin administration. Insulin was then administered directly into the portal vein. Following insulin injection, a second liver tissue sample was harvested from the left hepatic lobe at the designated time point. All collected liver tissues were immediately snap-frozen in liquid nitrogen to preserve protein and RNA integrity and subsequently stored at −80°C until further biochemical and molecular analyses.

### 2.11 *In-vitro* study

H4IIE cells were received from Dr. Young’s lab at the Vanderbilt University. Cells were maintained in DMEM low glucose medium. AAV8-TBG- (GFP or HuCETP) vectors were transfected into cells for experimental analysis. Cells were treated with or without BSA conjugated palmitate and testosterone overnight and then were incubated with 100nM of insulin for 15 min. Cells were collected and proteins were isolated for immunoblotting experiments.

### 2.12 Statistics

All the data are presented as mean and standard deviation (SD). Statistical analyses performed using GraphPad Prism, version 10.2.2. Significant differences within groups were analyzed by 1-way ANOVA with Bonferroni post-hoc comparisons or 2-way ANOVA with Tukey’s multiple comparison test as indicated in each figure legend. Repeated measures 1-way ANOVA with Bonferroni post-test comparison was used to statistically analyze blood glucose levels during OGTT. P-values <0.05 were considered statistically significant.

## 3. Results

### 3.1 AAV8-mediated expression of HuCETP in mouse liver

CETP is highly expressed in the human liver and secreted into circulation but is naturally absent in mice. To explore the physiological function of hepatic CETP, we utilized liver-directed expression of human CETP (HuCETP) to investigate whether dietary lipid excess regulates hepatic lipid metabolism and its link to liver disease progression in a sex- and CETP-dependent manner. Wild-type C57BL/6J mice of both sexes were injected with recombinant AAV encoding either GFP or HuCETP under the control of the liver-specific thyroxin-binding globulin (TBG) promoter, leading to predominant hepatic expression of GFP and HuCETP, respectively. AAV injection resulted in CETP expression found in circulation and liver, but not in other peripheral tissues (Fig. 1 A and B). Hepatic HuCETP (L-HuCETP) expression altered lipoprotein levels under chow-fed conditions (Fig. 1C-E). We found L-HuCETP reduced HDL-C and raised plasma TG consistent with findings from previous HuCETP transgenic mouse studies [21, 22]. Liver weight and lipid content were unchanged in chow-fed L-HuCETP mice compared with GFP controls (S.Fig.1 A-F). These data indicate that L-HuCETP expression has minimal effects on hepatic lipid metabolism under chow-fed diet conditions.

**Figure. 1.**
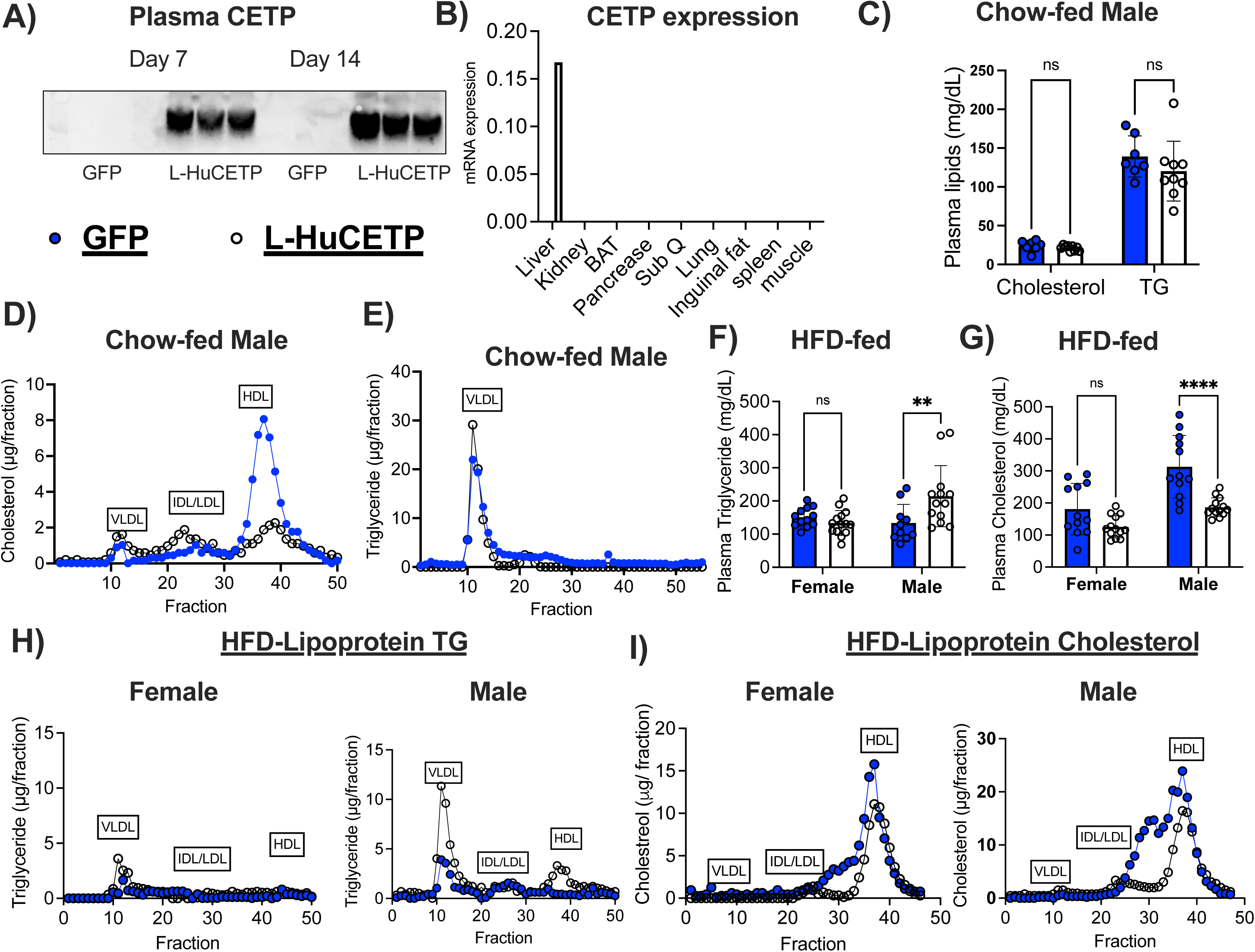
AAV8-mediated expression of HuCETP in mouse liver. A) CETP protein levels in plasma. B) Quantitative reverse transcriptase polymerase reaction(qRT-PCR) analysis of CETP in tissues. C) Quantification of fasting TG and cholesterol in serum from chow-fed mice. D-E) Cholesterol and TG distribution in lipoprotein in chow-fed mice. F-G) Fasting triglycerides and cholesterol in serum from HFD-fed mice. H-I) Lipid distribution in lipoprotein fractions from HFD-fed mice. Data are displayed as mean ± SD. Significant differences were determined using Student’s test A-E and two-way ANOVA used for F-J. The error bar indicates SD. *P<0.5; **P< 0.01; ns, not significant. GFP, green fluences protein; L-HuCETP, liver human cholesteryl ester transfer protein; TG, triglycerides; HDL-C, high-density lipoprotein cholesterol; VLDL-C, very low-density lipoprotein cholesterol.

### 3.2 Hepatic HuCETP expression altered lipoprotein lipid distribution and apolipoprotein levels in HFD-fed mice

In humans, the well-established function of CETP is to facilitate the net flux of cholesteryl esters from high-density lipoprotein (HDL) particles to low-density lipoprotein (LDL) and very-low-density lipoprotein (VLDL) particles, coupled with shuttling of triglycerides from LDL/VLDL to HDL particles. To determine the impact of L-HuCETP expression on diet-induced dyslipidemia, both male and female mice were fed a high-fat diet (HFD) for 15 weeks. In male mice, L-HuCETP significantly increased fasting triglyceride levels and lowered fasting cholesterol levels compared to GFP controls, whereas no differences in fasting lipid were observed in female mice (Fig F-G). Furthermore, L-HuCETP expression mice of both sexes exhibited higher VLDL-TG levels and lower HDL-cholesterol levels compared to GFP control mice (Fig.1 H-I). Additionally, we analyzed apolipoprotein levels in plasma by immunoblotting (S.Fig.2A) and observed that ApoC1 expression was significantly reduced in females, but no changes found in L-HuCETP male mice. In contrast, ApoB48 and ApoE levels were not changed in females but were elevated in males with L-HuCETP expression (S.Fig. 2 B-C). Collectively, these findings indicate that hepatic HuCETP expression alters lipoprotein lipid and apolipoprotein composition in a sex-specific manner following high-fat diet feeding.

**Figure. 2.**
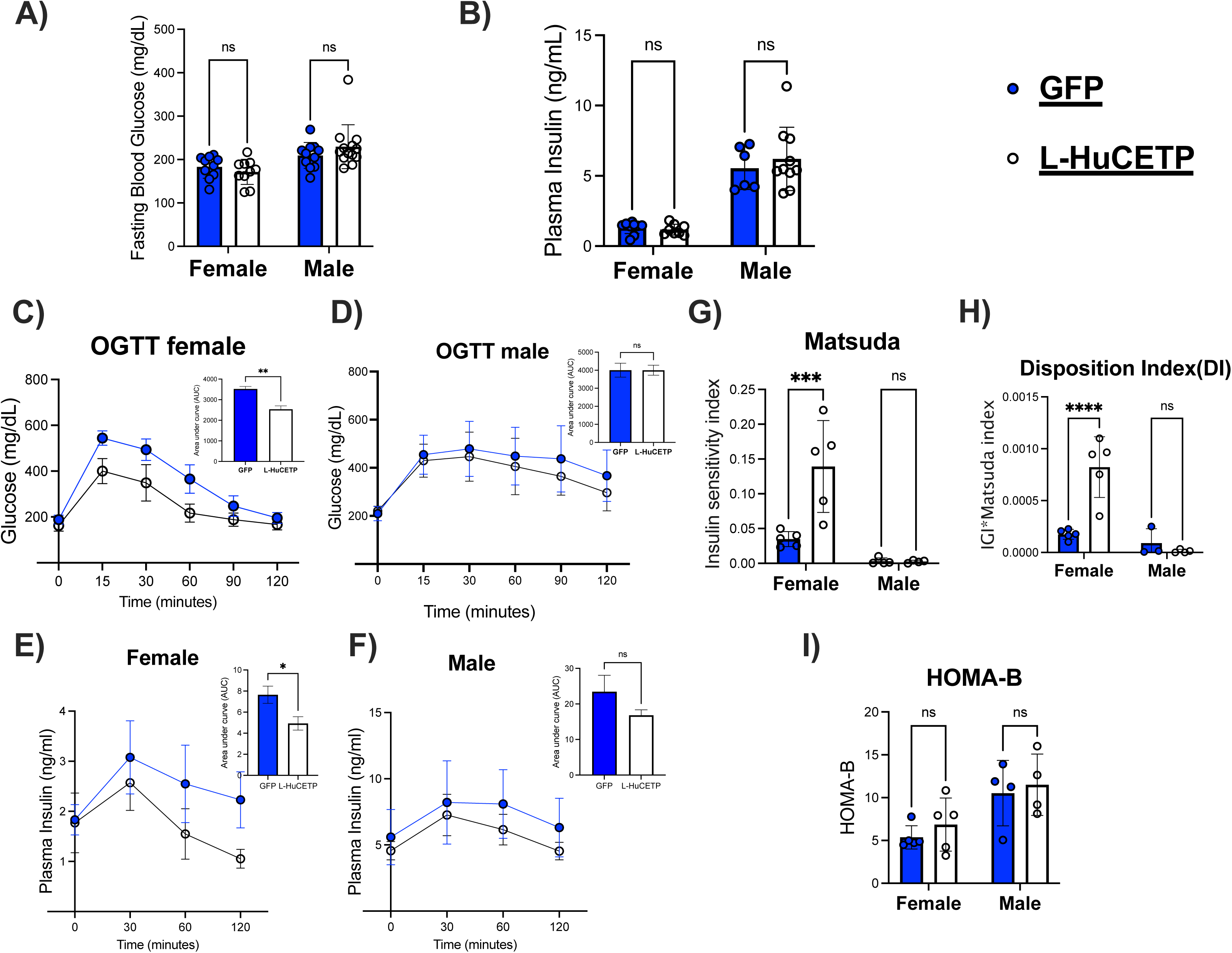
Hepatic HuCETP expression improves glucose tolerance and insulin sensitivity in females but not in males. Male and female GFP and L-HuCETP mice were fed a HFD for 8 weeks. A) Blood glucose B) Insulin levels were measured after 5 hours of fast condition. C) Oral glucose tolerance test and AUC in HFD fed female mice. D) Oral glucose tolerance test and AUC in HFD fed male mice. E) Insulin curve and AUC during oral glucose tolerance test in female. F) Insulin curve and AUC during oral tolerance test in male. G) Matsuda, H) Disposition Index (DI), I) HOMA-B were calculated from fasting glucose, insulin and glucose tolerance test values. Significant differences were determined by two-way ANOVA with Tukey’s multiple comparison post hoc analysis. The error bar indicates SD. *P<0.5; **P< 0.01; ns, not significant. GFP, green fluences protein; L-HuCETP, liver human cholesteryl ester transfer protein.

### 3.3 Hepatic HuCETP expression minimal effect on fat and lean mass ratio in response to high fat diet in both sexes

To determine whether L-HuCETP expression influences overall adiposity, body weight and composition were monitored biweekly in GFP and L-HuCETP mice of both sexes during HFD feeding. Growth curve analysis revealed no significant differences between GFP and L-HuCETP mice of either sex (S.Fig. 3A–B). Body composition analysis demonstrated comparable fat and lean mass between GFP and L-HuCETP groups in both sexes (S.Fig.3 C). These findings suggest that L-HuCETP expression does not significantly changes adiposity during HFD feeding.

**Figure. 3.**
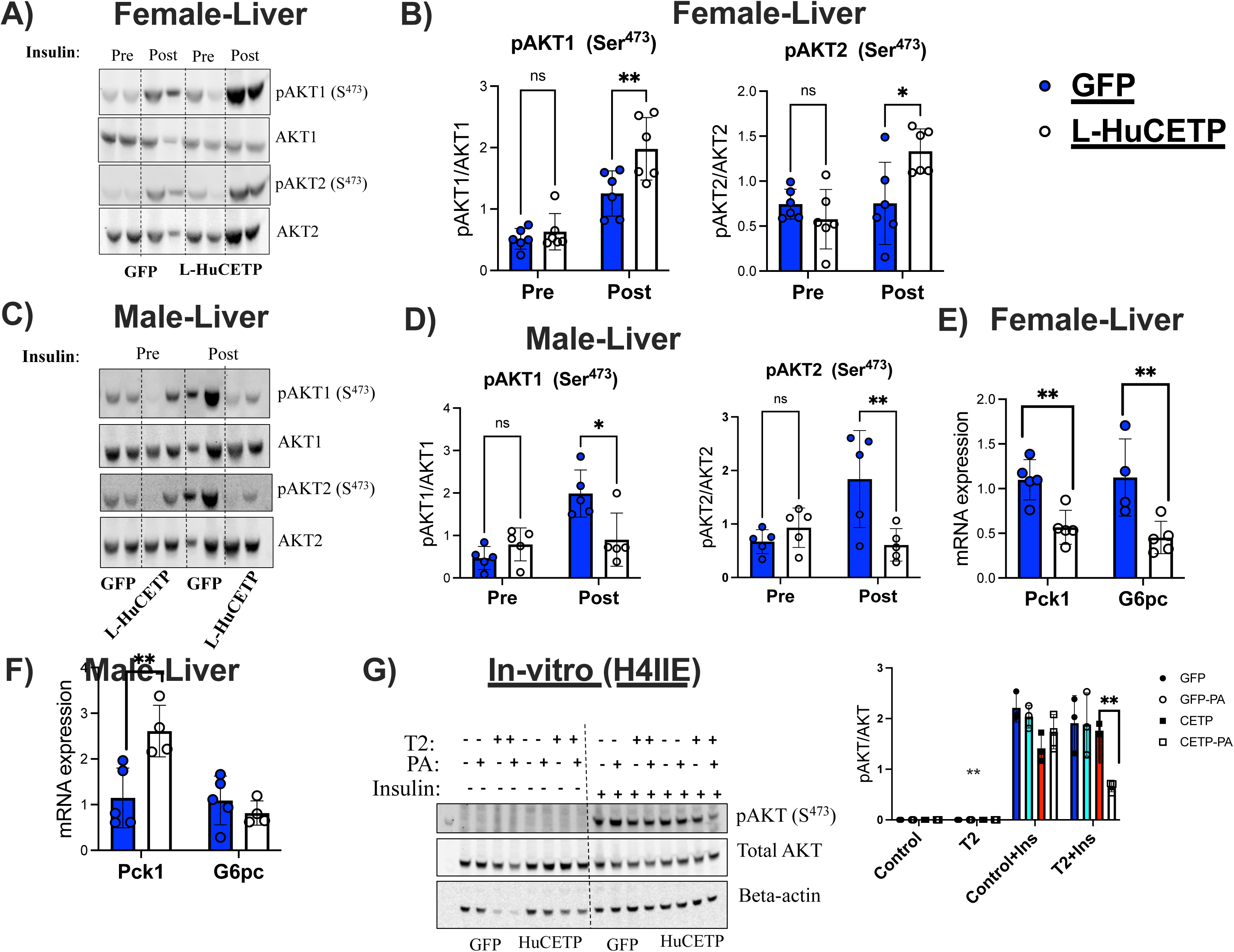
Hepatic HuCETP expression activates insulin-dependent AKT signaling in females but impairs this signaling in males. Female and Male Control (GFP) and L-HuCETP expression mice were fed a HFD (60%) for 15 weeks. A-B) Female, C-D) Male livers harvested from those mice Pre and Post insulin treatment for western blot of phopho-AKT1, phospho-AKT2, AKT1 and AKT2, and expression levels were determined by densitometry and normalized by respective total protein levels. E-F) liver gluconeogenesis genes Pck1 and G6pc mRNA expression in female and male respectively. G) H4IIE cells were treated with insulin and respective image of immunoblot of pAKT, total AKT and beta-actin and quantification data. Significant differences were determined by two-way ANOVA with Tukey’s multiple comparison post hoc analysis. The error bar indicates SD. *P<0.5; **P< 0.01; ns, not significant. GFP, green fluences protein; L-HuCETP, liver human cholesteryl ester transfer protein; PA-Palmitate; T2-testosterone; Ins-Insulin.

### 3.4 Hepatic HuCETP improves glucose tolerance and insulin sensitivity in female mice during HFD-feeding

We next examined whether L-HuCETP expression impacts glucose homeostasis and insulin action. Fasting blood glucose and insulin levels did not differ significantly between GFP and L-HuCETP mice in either sex, females had lower fasting insulin levels than males, regardless of CETP expression (Fig.2 A-B). We performed an oral glucose tolerance test (OGTT) after 8 weeks of high-fat diet feeding. Compared with HFD-fed GFP controls, HFD-fed L-HuCETP female mice displayed improved glucose tolerance, whereas male mice showed no significant difference (Fig.2 C-D). Furthermore, we measured insulin levels during OGTT. The results shown that female L-HuCETP mice exhibited lower insulin levels in the GTT compared with control, with no differences observed between male L-HuCETP and GFP mice (Fig.2 E-F). We further calculated insulin sensitivity and secretion indices from OGTT and plasma insulin data. Female L-HuCETP mice showed increased Matsuda index (a measure of insulin sensitivity during a GTT) and disposition index (DI, a measure of insulin secretion) relative to GFP controls, indicating improved insulin sensitivity and β-cell function. In contrast, male L-HuCETP mice displayed a no changes in Matsuda index, DI and HOMA-B (Fig.2 G-H). Taken together, these results indicate that hepatic-specific HuCETP expression enhances glucose tolerance and insulin sensitivity in female mice, whereas no such metabolic improvements were observed in male mice relative to their corresponding GFP control group.

### 3.5 Hepatic HuCETP expression improves hepatic insulin signaling in females but not in males

To better understand the mechanisms underlying the improved glucose tolerance in female L-HuCETP mice, we examined key insulin signaling pathways in the liver following portal vein insulin administration. In physiology, insulin secreted from pancreatic β-cells enters the circulation *via* the portal vein and first reaches the liver. To mimic this physiological route, under anesthesia fasted mice received insulin injection through the portal vein, and liver tissues were collected before and after insulin administration. In HFD-fed female mice, acute insulin treatment increased the phosphorylation of AKT at Ser473 in the liver in GFP controls, and expression of L-HuCETP further augmented insulin-stimulated AKT phosphorylation (Fig 3 B). Whereas in males, insulin stimulated pAKT-Ser473 in GFP controls, but expression of L-HuCETP blunted insulin-stimulated AKT phosphorylation (Fig 3 D). In females, L-HuCETP reduced levels of mRNA for gluconeogenesis genes glucose-6-phosphatase (G6pc) and phosphoenolpyruvate carboxykinase 1 (Pck1) (Fig.3 E). However, in males L-HuCETP increased Pck1 mRNA expression *vs.* control (Fig.3 F). Collectively, these observations suggest that improved hepatic insulin signaling may contribute to improved glucose tolerance by inhibiting hepatic glucose production in L-HuCETP mice, which is female specific.

To further explore mechanisms for the regulation of insulin signaling in L-HuCETP mice, we expressed HuCETP in H4IIE rat hepatoma cells and treated the cells with testosterone along with or without Palmitate (an *in vitro* model of hepatic steatosis and insulin resistant liver). H4IIE cells naturally lack CETP. Cells were transfected with AAV-GFP or AAV-HuCETP and treated with or without insulin in the presence or absence of testosterone. Immunoblot analysis revealed that HuCETP expression reduced insulin-stimulated AKT phosphorylation only when cells were treated with both testosterone and palmitate, suggesting androgens contribute to HuCETP-mediated impairment of insulin signaling (Fig.3 G). These *in vitro* findings corroborate our *in vivo* results, where male L-HuCETP mice failed to show enhanced insulin sensitivity compared with GFP controls. Collectively, these findings indicate that hepatic CETP expression enhances insulin sensitivity through activation of insulin signaling pathways in females, whereas in males testosterone may antagonize this effect.

### 3.6 Hepatic HuCETP expression worsens diet-induced hepatic steatosis in male mice

We next investigated the sex differences in liver lipid mechanism regulated by L-HuCETP during obesity. After 15 weeks of HFD-feeding, plasma free fatty acid (FFA) and liver triglyceride (TG) and levels were significantly elevated in male L-HuCETP mice compared with GFP controls, whereas no differences were observed in females across groups (Fig. 4A and B). Hepatic cholesterol levels were similar between L-huCETP and GFP mice in both sexes (Fig. 4C). Liver weight did not differ between L-HuCETP and GFP groups of either sex (Fig. 4D). Histological analysis by hematoxylin and eosin (H&E) staining revealed that male L-HuCETP mice exhibited more pronounced macro-steatosis than GFP controls, while no significant differences were detected between L-HuCETP and GFP groups in females (Fig. 4E). To understand hepatic lipid regulation, we next examined the expression of key proteins in pathways of lipogenesis, lipid uptake, and secretion. In males, L-HuCETP expression significantly increased the proteins of FASN, SCD1, ACC1 compared with GFP controls (Fig. 4F-G). Whereas in females CETP did not significant impacts these proteins did not significantly differ compared to controls. Interestingly, L-HuCETP expression did not alter LRP1 protein levels on male mice but significantly upregulated LRP1 protein levels on female L-HuCETP mice (Fig. 4F-G). Collectively, these results demonstrate a male-specific pathology that hepatic HuCETP expression promotes hepatic steatosis by increasing lipogenesis proteins in the liver.

**Figure. 4.**
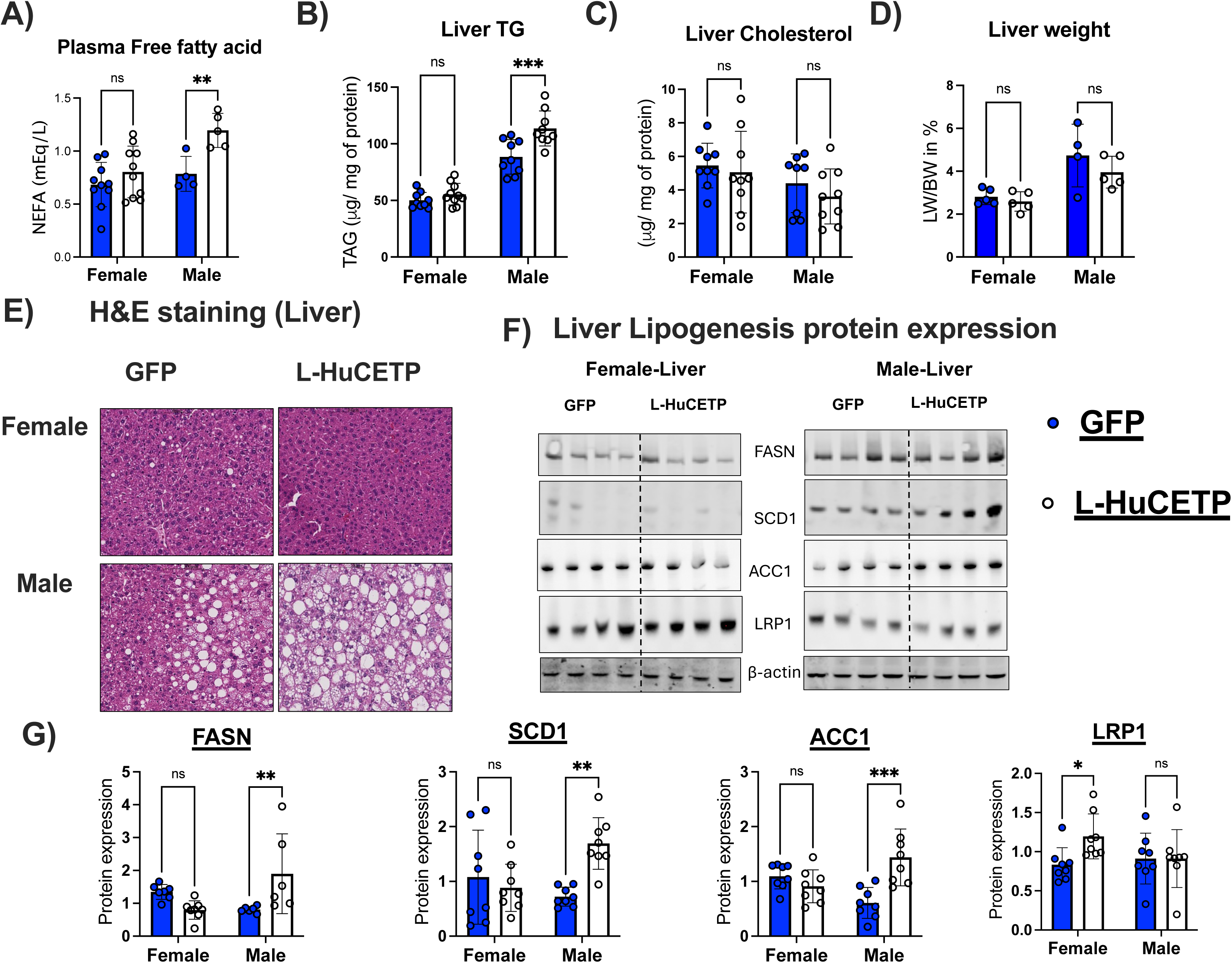
Diet-induced hepatic steatosis is worsened in male HuCETP mice. Mice were fed a HFD (60%) for 15 weeks. A) L-HuCETP increased fasting free fatty acid levels in males. B-C) Liver TG and cholesterol levels were measured. D) Liver weight E) Representative images of liver H&E staining. F-G) Representative western blot images for FASN, SCD1, ACC1 and LRP1, and the quantification results. Significant differences were determined by two-way ANOVA with Tukey’s multiple comparison post hoc analysis. The error bar indicates SD. *P<0.5; **P< 0.01; ns, not significant. GFP, green fluences protein; L-HuCETP, liver human cholesteryl ester transfer protein.

### 3.7 Hepatic expressions of HuCETP protects hepatic inflammation and early fibrosis in female mice but worsens in male mice

We next examined whether hepatic steatosis progressed to metabolic-associated steatohepatitis (MASH) in L-HuCETP mice. Plasma alanine aminotransferase (ALT) and aspartate aminotransferase (AST) were measured as markers of liver injury. Although ALT and AST levels were higher in males than females, no significant differences were detected between L-HuCETP and GFP controls in either sex, consistent with reports that liver enzymes do not reliably reflect fatty liver disease progression (S.Fig.4). In mouse models, the development of advanced liver fibrosis typically requires choline-deficient or GAN diet feeding, however prolonged HFD-feeding has also been reported to promote early stages of fibrotic progression. In the present study, mice were fed HFD for 15 weeks. Liver fibrosis was assessed by Masson’s trichrome staining, which detects collagen deposition as an indicator of early fibrosis. Female L-HuCETP mice displayed reduced early-stage fibrosis compared with GFP controls, whereas male mice showed increased early-stage fibrosis in the L-HuCETP group (Fig. 5A). To support these findings, we analyzed fibrosis- and inflammation-related gene expression by qRT-PCR. Expression of Tnfα, Ccl5, Acta2 and desmin genes were decreased in females, in line with histological results. Expression of Col1a was elevated in male L-HuCETP mice. (Fig. 5B-I). These findings suggest that L-HuCETP expression may modulate the early stages of pro-fibrotic and inflammatory responses in a sex-dependent manner. Specifically, L-HuCETP expression was associated with enhanced pro-fibrotic and inflammatory markers in male mice, whereas female mice exhibited a comparatively protective phenotype, characterized by attenuated fibrosis and inflammation.

**Figure. 5.**
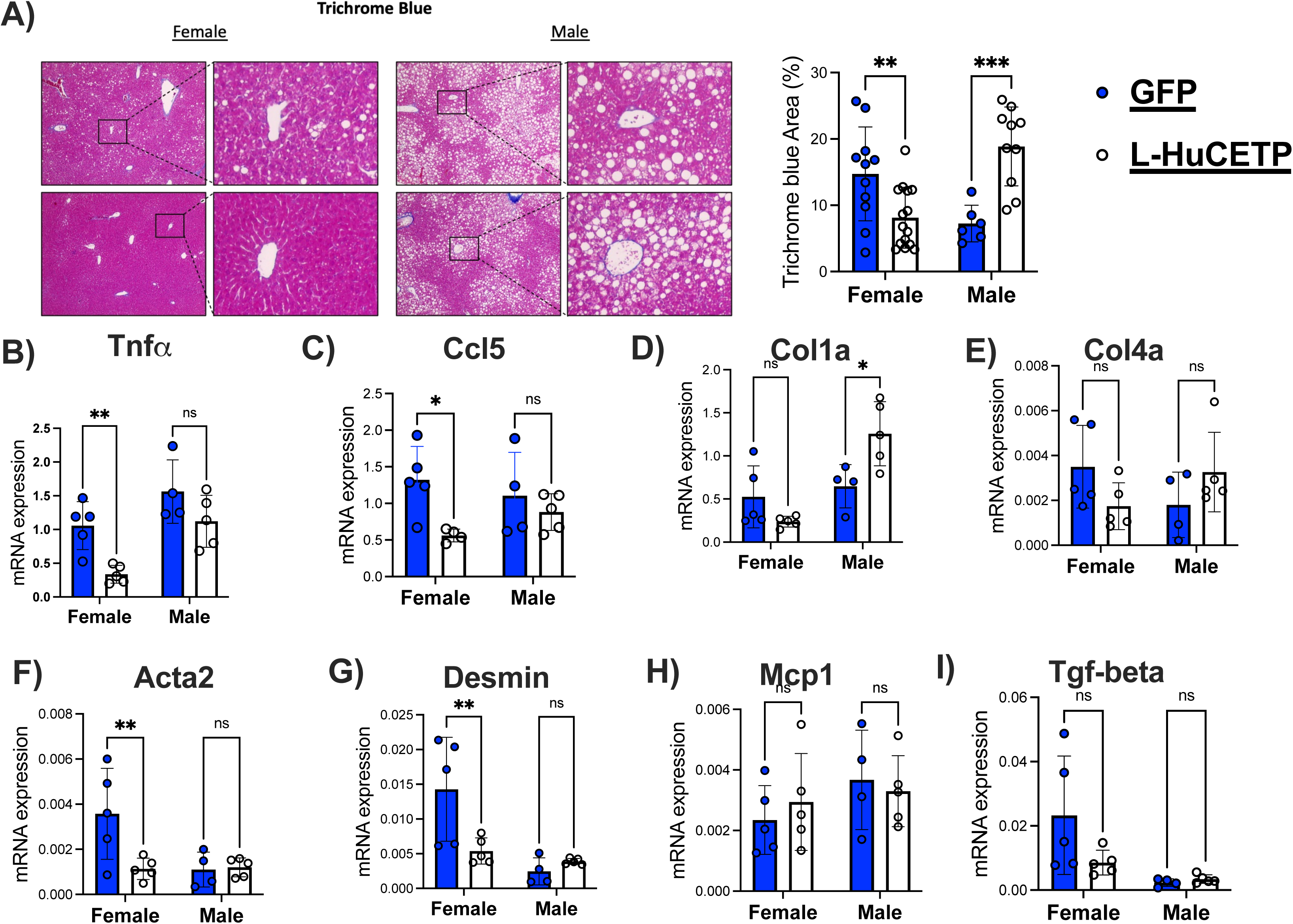
Hepatic HuCETP expression confers protection against diet-induced hepatic fibrosis and inflammation in a sex-dependent manner. Female and male GFP and L-HuCETP mice were fed a high-fat diet (60 %) for 15 weeks. A) Images of Trichrome blue staining and corresponding quantification data. B–H) Quantitative real-time PCR (qRT-PCR) analysis of mRNA expression levels of hepatic pro fibrosis-related genes (Col1a, Col4a, Acta2, and Desmin) and inflammation-related genes (Tnfα, Ccl5, Mcp1 and Tgf-beta). Significant differences were determined by two-way ANOVA with Tukey’s multiple comparison post hoc analysis. The error bar indicates SD. *P<0.5; **P< 0.01; ns, not significant. GFP, green fluences protein; L-HuCETP, liver human cholesteryl ester transfer protein.

### 3.8 Hepatic HuCETP expression controls metabolism in a sex-specific manner potentially involving ChREBP and HNF**α**

To define the molecular mechanisms underlying sex-dimorphism in hepatic metabolic regulation by HuCETP, we examined transcription factors governing gluconeogenic and lipogenic pathways in HFD-fed GFP and L-HuCETP mice. We analyzed the expression and subcellular distribution of ChREBP, HNF4α, and FOXO1 in female and male livers using nuclear and cytosolic fractionation. In females, L-HuCETP reduced nuclear ChREBP protein levels compared with GFP controls (Fig. 6A-B). In males L-HuCETP modestly increased cytosolic ChREBP levels to controls, but no change in nuclear abundance (Fig. 6C-D). Hepatic HNF4α expression was significantly reduced in male L-HuCETP mice, whereas no differences were observed in females (Fig. 6A-B). Additionally, cytosolic FOXO1 levels were significantly elevated in male HuCETP livers, with no changes in nuclear FOXO1 (Fig. 6C-D). Female mice showed no significant alterations in HNF4α or FOXO1 expression in either compartment (Fig. 6A-B). In males, altered cytosolic ChREBP and FOXO1 accumulated more with CETP, accompanied by reduced HNF4α, suggesting a distinct regulatory axis that may contribute to sex-specific metabolic outcomes.

**Figure 6.**
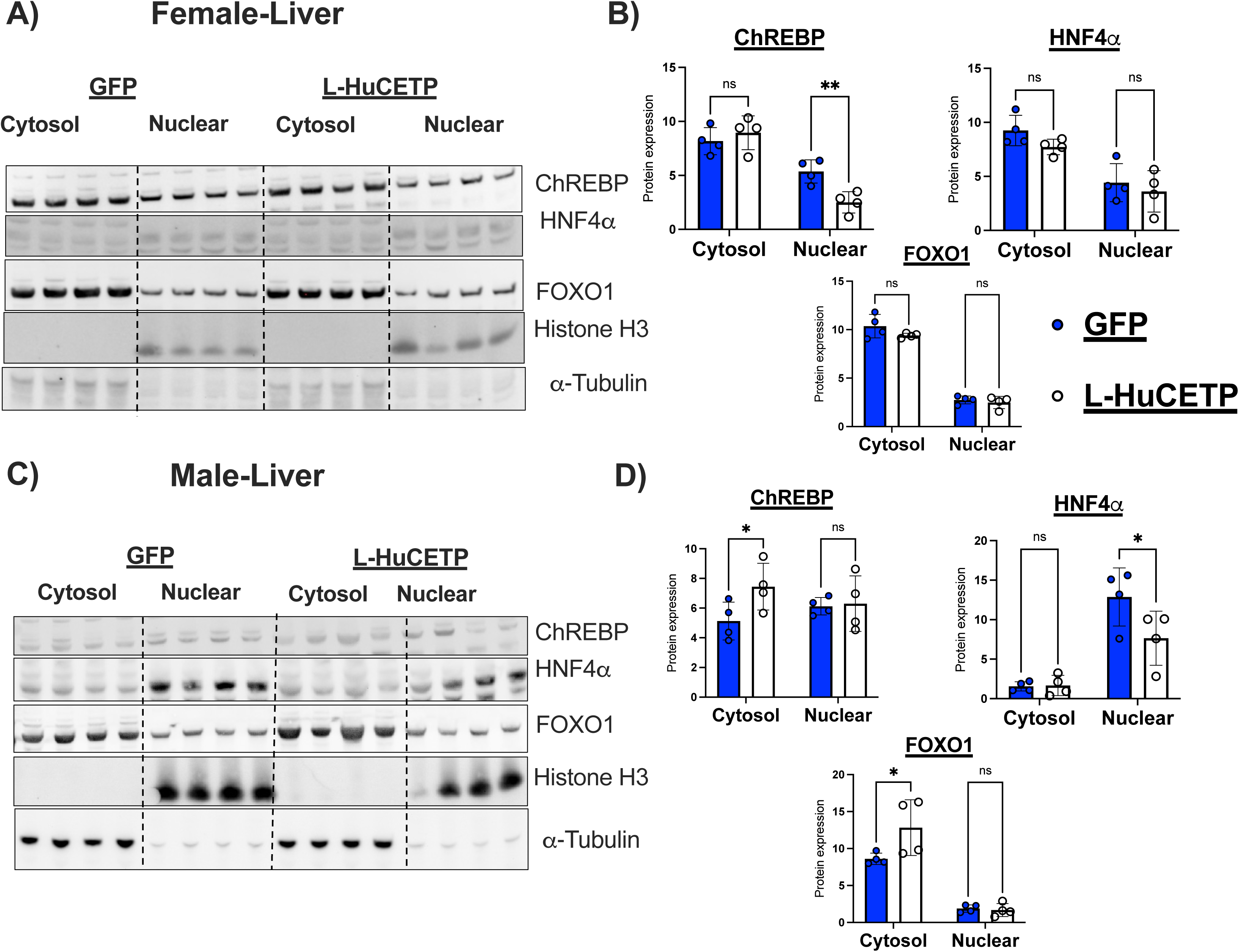
Hepatic HuCETP expression regulates ChREBP and HNF4_α_ subcellular location in a sex-dependent manner. Female and male GFP control and liver-specific HuCETP (L-HuCETP) mice were fed a high-fat diet (60 %) for 15 weeks. A–B) Representative immunoblot images and quantification of ChREBP, HNF4α, and FOXO1 protein expression in cytosolic and nuclear fractions of female liver. C–D) Representative immunoblot images and quantification of ChREBP, HNF4α, and FOXO1 protein expression in cytosolic and nuclear fractions of male liver. Histone H3 and α-tubulin were used as nuclear and cytosolic loading controls, respectively. Data are presented as mean ± SD. Statistical significance was determined by two-way ANOVA followed by Tukey’s multiple-comparison post hoc test. P < 0.05; P < 0.01; ns, not significant. GFP, green fluorescent protein; L-HuCETP, liver-specific human cholesteryl ester transfer protein.

### 3.9 Hepatic expression of HuCETP increases plasma estradiol levels in female and does not impact sex hormones in males

To understand the sex-dimorphism in liver metabolism, we determined CETP activity and its correlation to sex hormone levels in L-HuCETP mice. We found significant differences in circulating CETP activities between males and females (Fig. 7A). Circulatory estradiol levels were significantly elevated in female L-HuCETP mice compared with GFP controls, whereas no differences across phenotypes were detected in males (Fig. 7B). However, circulatory testosterone levels were not significantly altered in female and male L-HuCETP mice relative to controls (Fig. 7C). Furthermore, hepatic expression of ERα and androgen receptor (AR) was evaluated. ERα expression was significantly elevated in female HFD-fed L-HuCETP mice relative to GFP controls, whereas no significant changes were observed in male L-HuCETP mice. In contrast, AR expression remained unchanged in both female and male L-HuCETP mice compared with their respective control groups. Taken together, these findings suggest that elevated estradiol in female L-HuCETP mice may contribute to the protection against diet-induced insulin resistance, hepatic steatosis, and fibrosis progression. By contrast, the absence of such hormonal changes in males may underlie their increased susceptibility to metabolic dysfunction in the context of L-HuCETP expression.

**Figure. 7.**
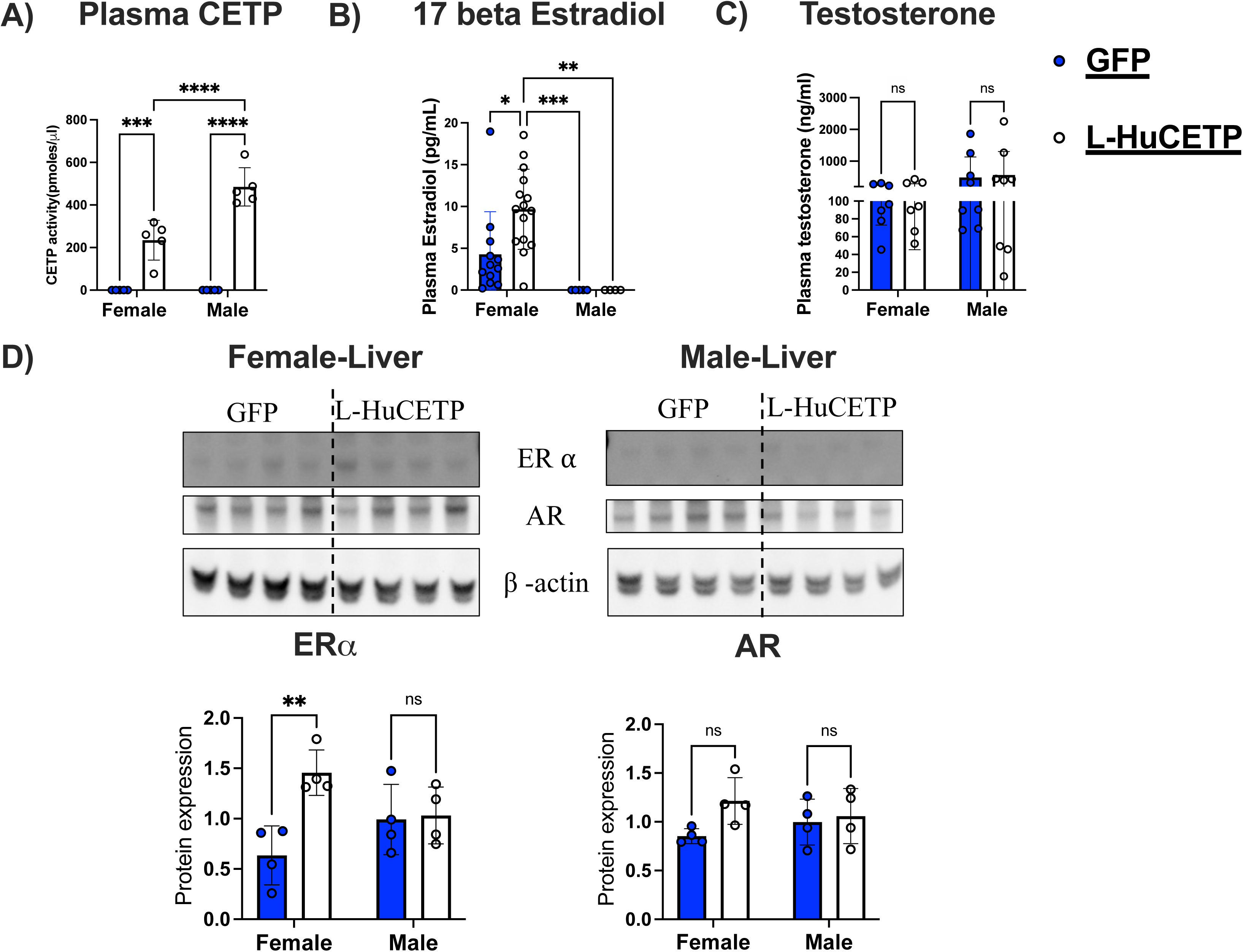
Hepatic-specific HuCETP expression is associated with increased estrogen levels in females. Female and male GFP and L-HuCETP mice were fed a high-fat diet (60 %) for 15 weeks. A) CETP activity B) Estradiol C) Testosterone measured from fasted mice serum. D) ERα and AR protein expression in liver. Significant differences were determined by two-way ANOVA with Tukey’s multiple comparison post hoc analysis. The error bar indicates SD. *P<0.5; **P< 0.01; ns, not significant. GFP, green fluences protein; L-HuCETP, liver human cholesteryl ester transfer protein.

## 4. Discussion

Numerous studies have demonstrated that females and males differ significantly in their physiology and metabolic homeostasis [24–26]. While obesity-induced insulin resistance and the progression of MASLD are well characterized, reliable data addressing sex-specific differences in dyslipidemia and MASLD remain limited. In this study, we demonstrate that hepatic expression of the human CETP gene differentially influences obesity, insulin resistance, and MASLD progression in a sex-dependent manner. Consistent with our previous CETP transgenic models, liver-specific HuCETP expression alters cholesterol distribution among circulating lipoproteins, supporting a conserved role for CETP in systemic lipid handling. Unlike our earlier work using whole-body HuCETP transgenic mice, here we employed a liver-directed approach using an AAV-TBG promoter, which allowed us to examine the acute effects of hepatic CETP and secreted CETP in the blood on fatty liver development and whole-body glucose metabolism. Given the central role of the liver in lipid metabolism-including lipoprotein uptake, assembly, and secretion-this model provides a more targeted framework to understand CETP function in hepatic physiology.

Beyond lipid metabolism, we found that L-HuCETP exerts several sex-specific effects on glucose homeostasis and its key regulatory mechanisms. In female mice, L-HuCETP expression significantly improved systemic glucose tolerance and enhances insulin signaling, whereas in male mice these benefits are absent. Instead, male mice exhibit impaired insulin signaling in response to L-HuCETP expression, indicating a sexually divergent metabolic response. Mechanistically, *in vitro* studies further support a role for sex hormones in mediating this divergence, as testosterone exposure selectively attenuated insulin signaling in HuCETP-expressing cells compared with controls. HuCETP may influence diet-induced fatty liver disease progression through sex-specific transcriptional programs. Our data implicate carbohydrate-responsive element–binding protein (ChREBP) as potential regulatory factor in females, whereas hepatocyte nuclear factor 4α (HNF4α) associates with HuCETP-driven hepatic responses in males. Further studies are needed to causally implicate these transcription factors in CETP’s sex-specific effects.

Our OGTT results demonstrate that diet-induced insulin resistance was significantly improved in female L-HuCETP mice compared with GFP controls by increasing pAKT, whereas no improvement in glucose tolerance was observed in male L-HuCETP mice. These results are consistent with previous studies using transgenic HuCETP models, which demonstrated that females are protected against diet-induced insulin resistance [20, 21]. Protection against insulin resistance was reduced in ovariectomized females, suggesting that sex hormones play a critical role in preventing obesity-induced metabolic dysfunction [27]. Clinical studies similarly report a higher prevalence of diabetes and its complications in obese men compared with obese women [28]. In contrast, menopause has been associated with impaired glucose homeostasis, while hormone replacement therapy improves glucose tolerance and reduces the risk of diabetes development [29, 30]. Together, clinical and experimental evidence clearly indicates that the development of insulin resistance in obesity is sex-dependent and disproportionately affects males [28]. Supporting this hypothesis, our *in vitro* studies demonstrated that HuCETP expression in H4IIE cells treated with testosterone resulted in downregulation of insulin signaling.

The current study demonstrates that hepatic expression of HuCETP differentially regulates the key gluconeogenic genes Pck1 and G6pc in a sex-dependent manner during the progression of HFD-induced liver disease. In normal physiology, insulin suppresses hepatic gluconeogenesis while promoting lipid synthesis; however, in insulin resistance, gluconeogenesis increases despite continued lipogenesis, a phenomenon known as selective insulin resistance[31, 32].

Our results support that hepatic lipid accumulation in L-HuCETP male mice is driven by increased hepatic lipogenesis, as evidenced by elevated expression of key lipogenic enzymes, including ACC1, FASN, and SCD1. Studies have shown that diet-induced obesity increases plasma FFA in both females and males due to enhanced adipose tissue lipolysis [33]. However, females relatively protected against FFA-induced insulin resistance [34, 35]. Elevated plasma FFA levels in L-HuCETP males were associated with the development of hepatic steatosis shown by H&E staining and increased hepatic triacylglycerol (TAG) accumulation. Numerous studies have reported sex-specific differences in the development of diet-induced fatty liver disease, with outcomes varying depending on diet composition and mouse strain [36–39]. Both experimental and epidemiological studies have demonstrated that hepatic steatosis is more prevalent in males than females [40]. Our study we observed that L-HuCETP appears may be protective in females, but in males it is associated with increased pro-inflammatory and pro-fibrosis gene expression. Future study will focus on determine the role of CETP on sex specific liver inflammation and fibrosis progression.

Liver-specific inhibition of the *Mlxipl* gene, which encodes ChREBP, suppresses lipogenesis and alleviates fatty liver in obese mice [41]. In the present study, we observed lower ChREBP expression in female HuCETP mice, which may contribute to the protection against hepatic lipid accumulation seen. HNF4α, a key hepatic nuclear receptor, is downregulated in NAFLD and metabolic disease models, and its loss is known to reduce plasma lipids while causing steatosis through defective VLDL secretion [42, 43]. Conversely, HNF4α overexpression protects against diet-induced steatosis [44]. Interestingly, we observed significantly higher circulating estrogen levels and increased hepatic ERα expression in female L-HuCETP mice. These findings suggest that the protective effects of CETP observed in female mice may be mediated, at least in part, through estrogen signaling pathways rather than androgen-dependent mechanisms. However, the precise molecular mechanisms linking CETP expression to estrogen signaling remain unclear. Future studies are warranted to elucidate the interaction between CETP and ERα-mediated pathways and to determine whether modulation of estrogen signaling directly contributes to the sex-specific protective effects associated with CETP expression.

In summary, L-HuCETP regulates diet-induced insulin resistance and hepatic steatosis in a sex-dependent manner. In male mice, androgens per se appear to exert beneficial metabolic effects, including in the liver (as seen in Liver KO AR models), raising the question of why human males are at greater metabolic risk. Notably, humans express CETP, whereas mice naturally lack it. Our findings suggest that restoring hepatic CETP amplifies beneficial metabolic effects in females, while in males it induces a gain-of-function interaction with androgens that may contribute to the increased risk of MASLD. Together, our results identify a sex-specific role for CETP in coordinating metabolic pathways and highlight its potential as a therapeutic target in metabolic disease.

## Author Contributions

S.C. and J.S. developed the overall concept and study design. S.C, U.A and LZ performed the experiments. S.C. and J.S. analyzed the data and prepared the figures. S.C. wrote the original draft of the manuscript; L.Z. and J.S. made the editing. All authors approved the final version

## Acknowledgements

The authors acknowledge the assistance of the Vanderbilt Mouse Metabolic Phenotyping Core (supported by NIH grant DK59637) and the support of Vanderbilt Transitional Pathology Shared Resource Core (supported by NCI/NIH Cancer Center Support Grant P30CA068485).

## Abbreviations

MASLD: Metabolic-Associated Steatotic Liver Disease
HuCETP: Human cholesteryl ester transfer protein
GFP: Green Fluorescence Protein
HFD: High fat diet
ChREBP: Carbohydrate-responsive element-binding protein
VLDL: Very Low-Density Lipoprotein
HDL: High Density Lipoprotein LDL-Low Density Lipoprotein
ApoB: Apolipoprotein B
AAV: Adeno-associated virus
TBG: Thyroxine-binding globulin
FASN: Fatty acid synthase
ACC1: Acetyl-CoA carboxylase 1
SCD1: Stearoyl-CoA desaturase-1
LRP1: Low density lipoprotein receptor-related protein 1
HNF4α: Hepatocyte nuclear factor 4α
FOXO1: Forkhead box protein O1
Pck1: Phosphoenolpyruvate carboxykinase 1
G6pc: Glucose-6-Phosphatase Catalytic Subunit 1
FFA: Free fatty acid
DNL: De novo lipogenesis

---

**Supplementary Figure 1.**
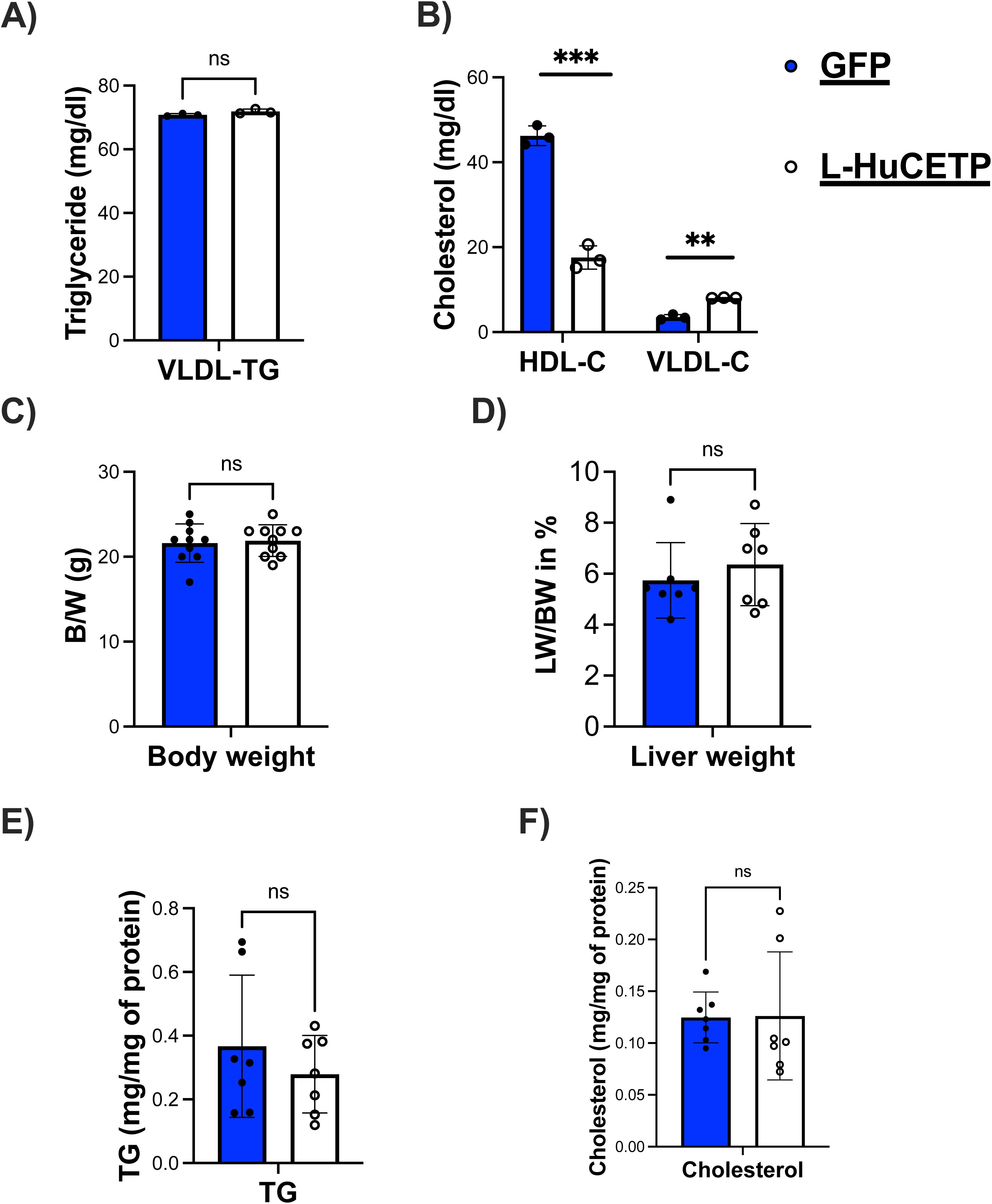
AAV-mediated liver-specific HuCETP expression does not alter plasma lipid levels or body weight. Wild-type mice were injected with AAV-GFP or AAV–liver-specific HuCETP (L-HuCETP) vectors and maintained on a chow diet. A-B) Plasma VLDL–triglyceride (TG) and HDL–cholesterol levels. C-D) Body weight and liver weight. E-F) Hepatic triacylglycerol (TG) and cholesterol content. Data are presented as mean ± SD. Statistical significance was determined as indicated: P < 0.05; P < 0.01; ns, not significant. GFP, green fluorescent protein; L-HuCETP, liver-specific human cholesteryl ester transfer protein.

**Supplementary Figure 2.**
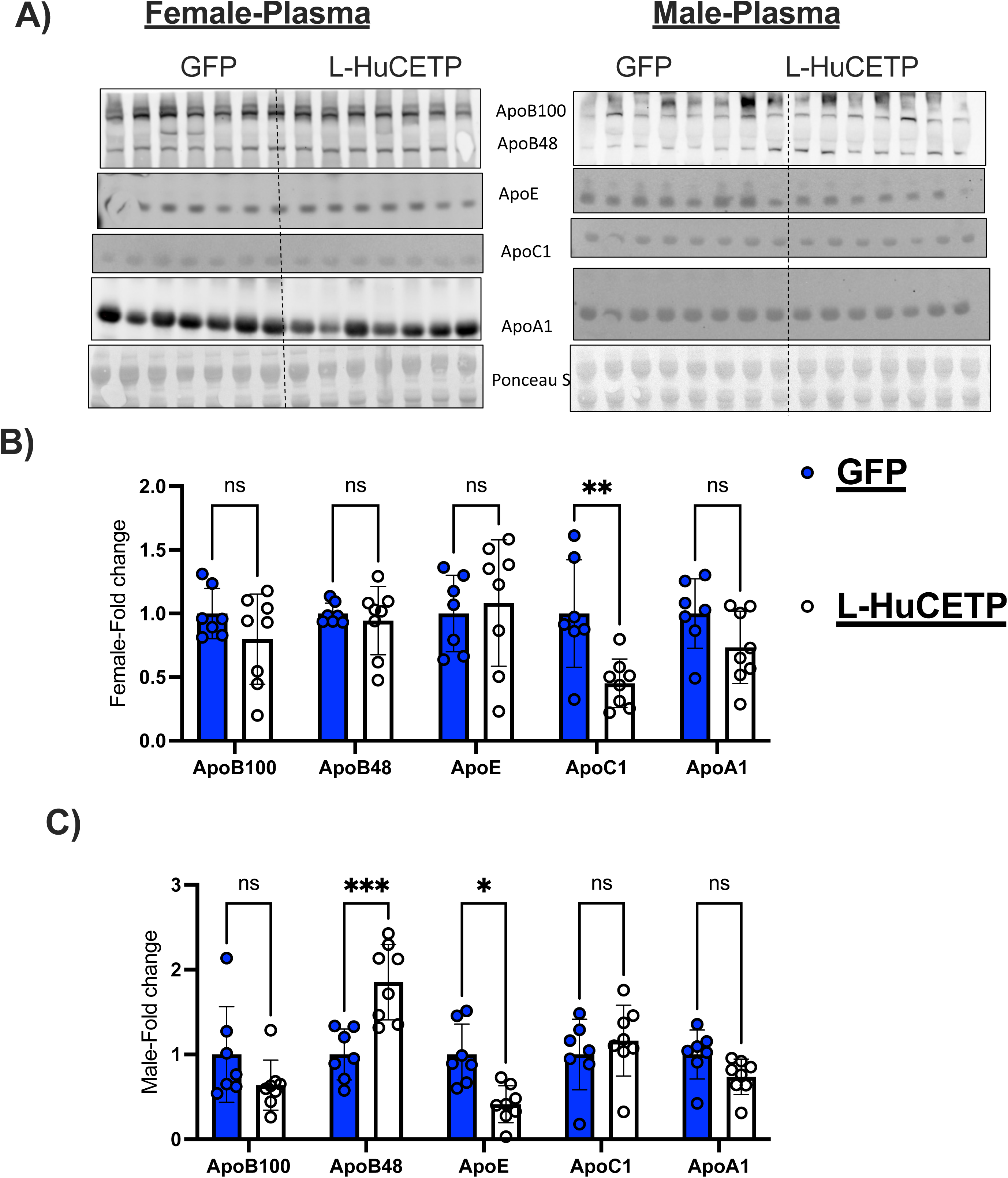
Hepatic HuCETP expression alters plasma lipoprotein profiles in a sex-dependent manner. Female and male GFP control and liver-specific HuCETP (L-HuCETP) mice were fed a high-fat diet (60 %) for 15 weeks. A) Representatives immunoblot images of plasma lipoproteins, including ApoB100, ApoB48, ApoE, ApoC1, and ApoA1, from female and male mice. B) Quantification of plasma lipoprotein protein levels in female mice. C) Quantification of plasma lipoprotein protein levels in male mice. Data are presented as mean ± SD. Statistical significance was determined as indicated: P < 0.05; P < 0.01; ns, not significant. GFP, green fluorescent protein; L-HuCETP, liver-specific human cholesteryl ester transfer protein.

**Supplementary Figure. 3.**
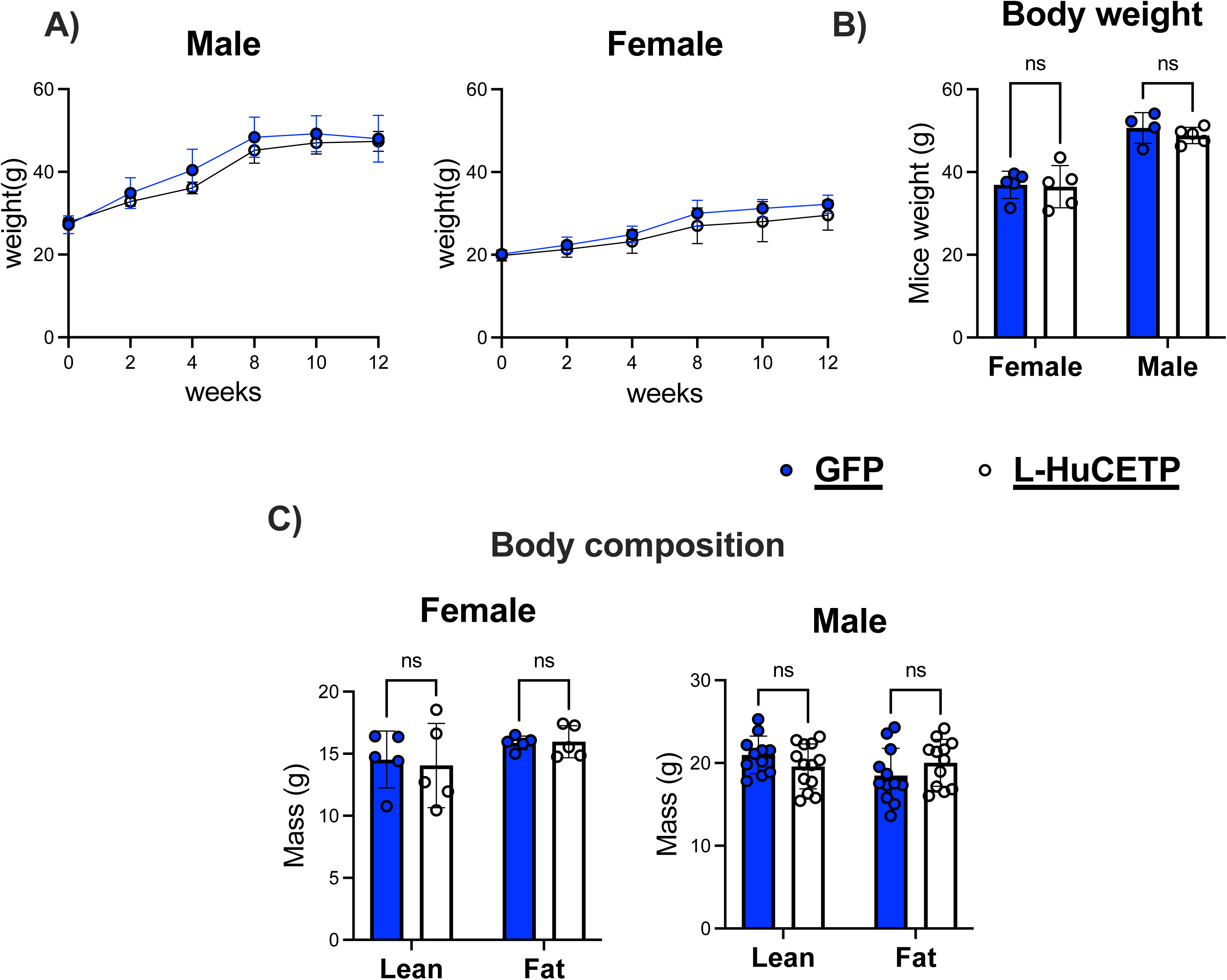
Liver expression of HuCETP has no effect on body weight in HFD feeding. GFP and L-HuCETP Female and male mice were fed a HFD (60%) for 15 weeks. A) Body weights were measured every 2 weeks. B) Whole body weights were measured, C) Body lean and fat mass were recorded. Significant determined by two-way ANOVA with Tukey’s multiple comparsion post hoc analysis. The error bar indicates SD. *P<0.5; **P< 0.01; ns, not significant. GFP, green fluences protein; L-HuCETP, liver human cholesteryl ester transfer protein.

**Supplementary Figure 4.**
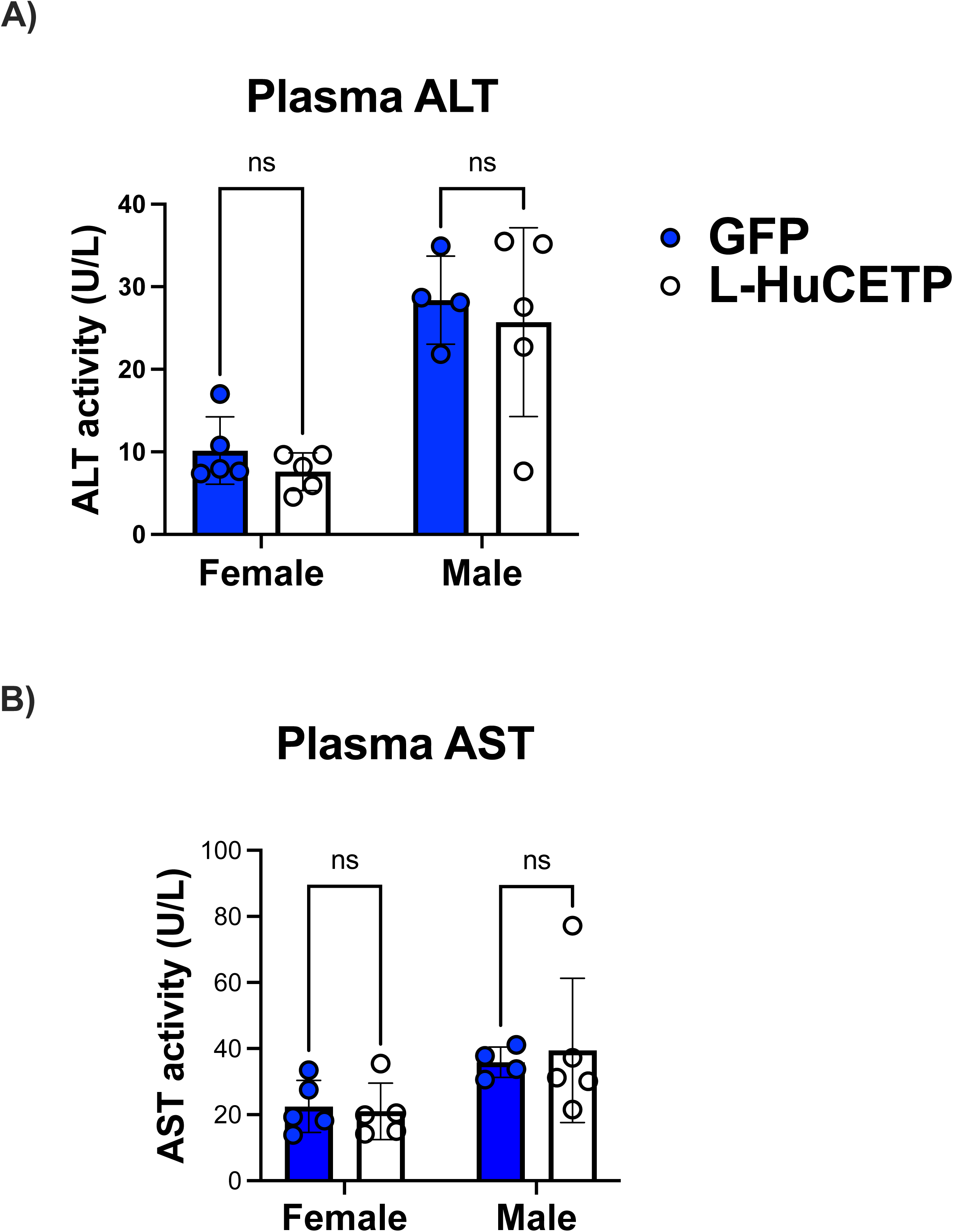
Effect of liver-specific HuCETP expression on plasma liver enzyme activity. Female and male GFP control and liver-specific HuCETP (L-HuCETP) mice were fed a high-fat diet (HFD; 60% kcal from fat) for 15 weeks. A) Plasma alanine aminotransferase (ALT) activity. B) Plasma aspartate aminotransferase (AST) activity.

